# Ebola virus mRNAs contain RNA structures that are critical for viral infection and targetable by antisense oligonucleotides

**DOI:** 10.64898/2026.09.04.749507

**Authors:** Michelle Luo, Judith Olejnik, Kristina Meier, Tanja Hann, Elke Mühlberger, Anna Marie Pyle

## Abstract

Filoviruses, such as Ebola virus (EBOV), are highly pathogenic non-segmented negative-sense RNA viruses (nsNSVs) with limited therapeutic options. Filovirus RNA structures remain largely untapped due to the enhanced biosafety requirements for handling infectious virus. Here, we present the first in-cell secondary structure maps of four EBOV mRNAs (VP35, VP40, VP30, and VP24) using two orthogonal chemical probing approaches: SHAPE-MaP and fbDMS-MaP. We find that EBOV mRNA coding sequences (CDS) are highly structured, much like +ssRNA viruses, whereas untranslated regions (UTRs) are significantly less structured. This suggests that high CDS structure contents are general features of viral translation templates, and that nsNSVs have evolved separate regulatory function at the RNA structure level that extends beyond using distinct mRNAs and genomes. These structure maps are consistent with formation of mRNA 5′ hairpin structures during infection and reveal numerous additional RNA structures within the CDS, 3′ UTRs, and at CDS-UTR junctions. To assess functionality, we disrupted these structures with locked nucleic acid (LNA) antisense oligonucleotides. Disrupting the TSS hairpins in VP35, VP30, and VP24 decreased infection by >60%, indicating these mRNA structures are critical for infection. LNA targeting of the newly identified structures reduced EBOV infection by 31% to 88%, thereby linking RNA structural integrity to viral function. Synonymous mutation rates and covariation analysis provided evolutionary support across mammalian filoviruses for the functional RNA elements observed. Collectively, these results demonstrate EBOV mRNAs contain numerous conserved RNA motifs contributing to viral infection, and that these elements represent promising targets for development of pan-filoviral therapeutics.

**Importance:** EBOV and related filoviruses pose a significant global health threat, yet our understanding of the RNA architectural mechanisms driving infection remains incomplete. Due to enhanced biosafety constraints, filovirus RNA structures have not been mapped in cells. Here, we experimentally map secondary structures of four EBOV mRNAs at biosafety level 4. We find that structure content is concentrated within protein coding regions, resembling the highly structured genomes of positive-sense RNA viruses, while untranslated regions are relatively unstructured. This separation of relative structural content reflects how translation and replication are delegated between mRNAs and genomes in negative-sense RNA viruses. We also show that these structures are critical for infection, with disruption reducing infection more than 80% in liver cells. Evolutionary analyses suggest a set of these regulatory elements are conserved across mammalian filoviruses. Broadly, this work expands the repertoire of filovirus regulatory elements, revealing new potential targets for developing pan-filoviral therapeutics.

## Introduction

Ebola virus (EBOV) is a member of the filovirus family, a group of non-segmented negative-sense RNA viruses (nsNSVs) ranking among the most virulent human pathogens (1). Filoviruses pose a major threat to global public health, causing severe hemorrhagic fevers with case fatality rates reaching up to 90% (2, 3). Since research on these pathogens requires biosafety level 4 (BSL-4) containment, this limits experimental accessibility and hinders our mechanistic understanding of filovirus infection. Consequently, treatment options for filovirus diseases are sparse, with only two licensed vaccines and two monoclonal antibody treatments against EBOV (4, 5). These countermeasures are ineffective against other filoviruses, which hinders response efforts during outbreaks of rarer filoviruses, like the current Bundibugyo virus outbreak in the Democratic Republic of the Congo and Uganda (6). Together, these challenges underscore the urgent need to identify conserved targets for pan-filoviral therapeutic development.

Since EBOV is the most extensively characterized filovirus, it is an ideal model system for studying filovirus biology. Its ∼19 kb negative-sense RNA genome is replicated through a positive-sense antigenome that serves as the template for genome synthesis. During transcription, seven positive-sense mRNAs encoding nine proteins are produced (**Fig 1A**) (7). These include the nucleoprotein (NP), RNA-dependent RNA polymerase (L), glycoprotein (GP), soluble– and small-soluble glycoproteins (sGP and ssGP), and viral proteins VP35, VP40, VP30, and VP24 that are involved in essential processes like RNA polymerase activity, transcription regulation, virion assembly, budding, and host immunomodulation (8). These proteins are encoded within a diverse RNA landscape that requires careful regulation for proper gene expression and viral infection.

**Fig 1.**
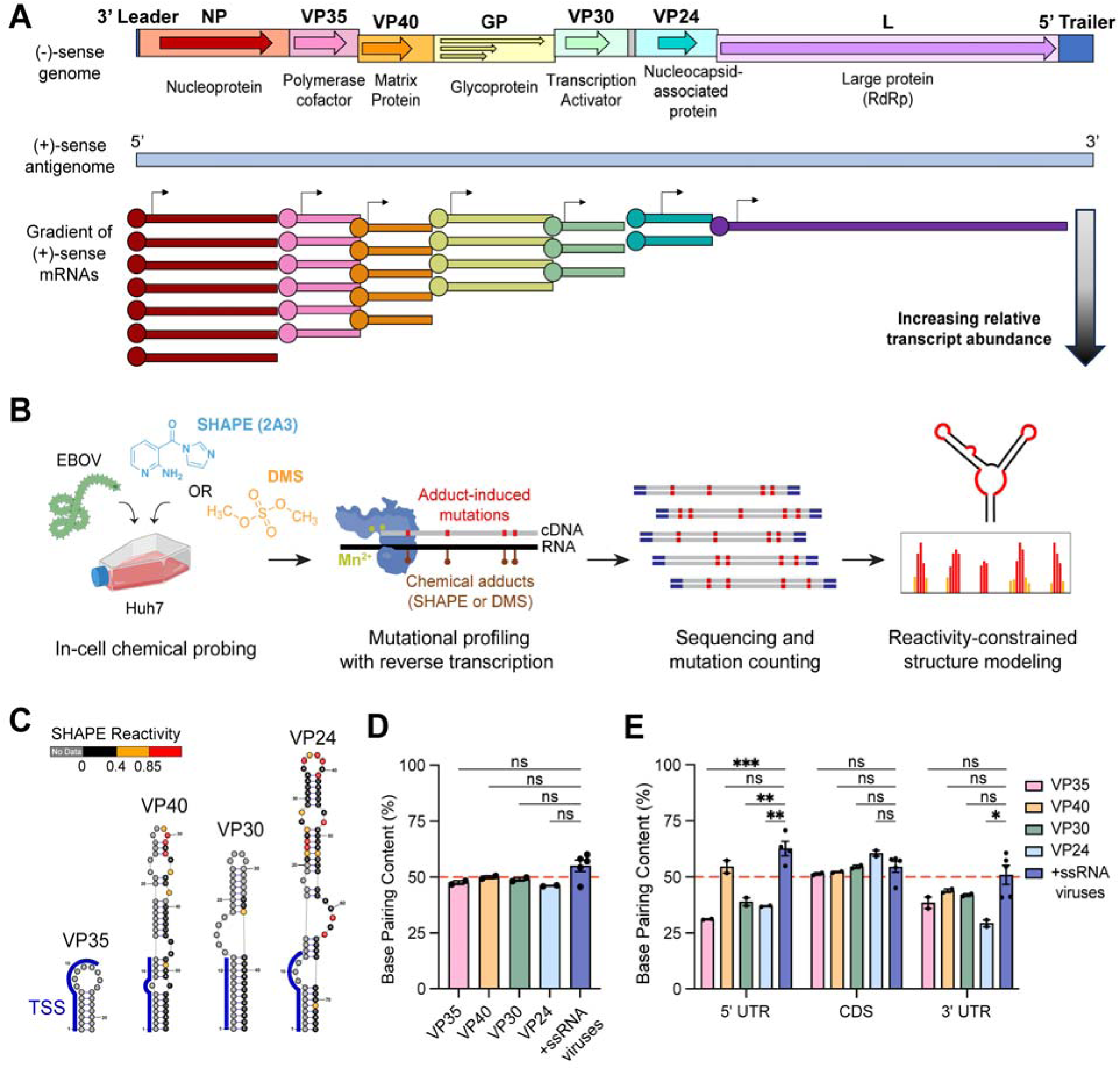
Structural architecture of EBOV mRNAs determined by chemical probing. (**A**) EBOV RNA landscape consisting of the (top) 19 kb genome with coding sequences indicated by arrows, (middle) 19 kb antigenome, and (bottom) transcription gradient of capped mRNAs. Genome features are to scale. mRNA transcripts are not to scale. (**B**) Schematic of in-cell chemical probing and structure prediction workflow. Infection schematic (left) partially created with Biorender. (**C**) mRNA 5’ hairpins containing the transcription start signal (TSS, blue) in the SHAPE-constrained structure maps. (**D-E**) Base-paired nucleotides of (**D**) full-length mRNAs or (**E**) specific mRNA regions. Data points represent BPC ± SEM of individual SHAPE replicates or comparable in-cell SHAPE studies of +ssRNA viral genomes: DENV2 (11); SARS-CoV-2 (12); HCV (13); and WNV (14). Red dashed line: BPC = 50%. Statistical significance: * p < 0.05; ** p < 0.01; *** p < 0.001; by ordinary one-way ANOVA for multiple comparisons.

Recent studies have shown that RNA viruses have elaborate RNA architectures that control key steps in viral infection, including genome replication, translation, and evasion of host defenses (9). These conserved RNA structures represent viable drug targets, underscoring the importance of studying viral RNA structures under physiological infection conditions (10). In the past decade, in-cell chemical probing has revolutionized the ability to sensitively map functional secondary structures in positive-sense single-stranded RNA (+ssRNA) viruses, such as Dengue virus (DENV), SARS-CoV-2, hepatitis C virus (HCV), West Nile virus (WNV) (11–14). For other types of viruses, such as segmented negative-sense viruses and retroviruses, structure mapping has been limited to influenza A virus and human immunodeficiency virus, while the RNA regulatory landscape of filoviruses and other nsNSVs is relatively uncharacterized (15–18).

RNA structures have been implicated in the regulatory processes of nsNSVs, but comprehensive experimental characterization of nsNSV architecture has been limited (19–25). Previous studies have focused on hairpin structures in the noncoding regions of filovirus genomes and the analogous stem loops in the extreme 5’ termini of the mRNAs (22, 23). These hairpin structures harbor the 12-nt transcription start sequence (TSS), both of which are critical for VP30-dependent transcription (22, 26). Another study showed a small stem loop in the genomic trailer interacts with the host chaperone HSPA8 to facilitate minigenome replication (27). However, little is known about filoviral RNA architectures outside of these short non-coding regions. As a rare exception, a stable hairpin structure has been shown to regulate RNA editing in the GP gene, affecting subsequent expression levels of GP, sGP, and ssGP, which control host entry and viral pathogenicity (28–30). While these studies highlight the importance of RNA structures, they rely on simplified systems like *in silico* predictions or minigenome replicons that do not recapitulate critical steps in viral infection, such as cell entry, virion assembly, or host immunosuppression (31, 32). A recent chemical probing study demonstrated that disrupting host-virus RNA-RNA interactions in respiratory syncytial virus (RSV) reduced infection in human lung cells, but the viral RNA structures were not comprehensively characterized (33). As a result, it remains unclear how extensively structure-based regulation occurs throughout the genomes or mRNAs of nsNSVs, how structures in coding transcripts could regulate viral infection, and whether these structures represent viable drug targets in filoviruses.

In this study, we use two orthogonal in-cell chemical probing methods at BSL-4 to comprehensively map the RNA secondary structures of four EBOV mRNAs (VP35, VP40, VP30, and VP24). We identified structured elements throughout the mRNAs, including the coding sequences (CDS), untranslated regions (UTRs), and CDS-UTR junctions. We compared the base-pairing contents of EBOV mRNAs to those of highly structured +ssRNA virus genomes (e.g. coronaviruses, flaviviruses) and observed comparable structure content within CDS regions. Disrupting the previously predicted TSS hairpins and newly identified elements with antisense oligonucleotides demonstrated these mRNA secondary structures are critical for EBOV infection. Synonymous mutation rates and covariation provided evolutionary support for a set of these functional RNA elements across mammalian filoviruses. Together, this study describes the first systematic characterization of RNA structures in a filovirus at BSL-4 and demonstrates that conserved RNA secondary structures may provide a tractable route to a pan-filoviral therapeutic.

## Results

### In-cell chemical probing of EBOV mRNAs under BSL-4 conditions

To elucidate the structural landscape of EBOV mRNAs in infected cells, we applied chemical probing and mutational profiling (MaP) to four EBOV mRNAs: VP35, VP40, VP30, and VP24. We tested two orthogonal probing methods: SHAPE-MaP (<u>S</u>elective 2’-<u>H</u>ydroxyl <u>A</u>cylation analyzed by <u>P</u>rimer <u>E</u>xtension) and fbDMS-MaP (four-<u>b</u>ase <u>Dim</u>ethyl <u>S</u>ulfate) (34–36). SHAPE probes, such as 2-aminopyridine-3-carboxylic acid imidazolide (2A3), selectively react with the 2’OH of flexible nucleotides to covalently deposit a bulky aromatic adduct (37). In contrast, DMS methylates the Watson-Crick faces of unpaired nucleotides, including uridine and guanine in an alkaline environment (35). These chemical adducts induce cDNA mutations during reverse transcription with a non-native cofactor (e.g. Mn^2+^) (38, 39). To capture the mutation patterns on an entire mRNA in a single pass, we utilized the ultra-processive reverse transcriptase, MarathonRT (39, 40). Deep sequencing identifies these mutational signatures to generate a reactivity profile, which is used to constrain secondary structure predictions (**Fig 1B**) (34).

All sequencing data passed the ShapeMapper2 quality controls, meeting the threshold read depths (>5000 read depth), mutation rates above background (>50%), minimal background mutation rates (<5%), and highly reactive nucleotides (>8%) (41). Mutation rates in chemically modified samples were significantly higher than untreated controls for both SHAPE and fbDMS datasets, indicating the RNAs were successfully modified (**Fig S1A-D, S1M-P**).

To assess reproducibility, we analyzed the correlation of normalized reactivities from two independent replicates of in-cell SHAPE-MaP and fbDMS-MaP. Pearson correlation coefficients (r) exceeded 0.85 for all SHAPE datasets and >0.80 for all fbDMS datasets (r>0.90 excluding G nucleotides) (**Fig S1E-H, S1Q-T**). Correlation values were also highly comparable between 2A3 and DMS probes (Δr <0.07), indicating that nsNSVs are amenable to in-cell structure probing with either reagent. For both probes, Pearson correlations decreased sequentially with transcript abundance, where (r_VP35_ > r_VP40_ > r_VP30_ > r_VP24_), suggesting that reactivity correlation and data quality scale with transcript copy numbers. Nonetheless, all datasets had correlations of r >0.80, which matches or exceeds those observed in +ssRNA virus probing studies (12–14).

### Structure content of EBOV mRNAs is higher in the coding sequences than the UTRs

To generate mRNA structure maps, we used the SHAPE reactivities to constrain SuperFold secondary structure predictions (38). Mapping SHAPE and fbDMS reactivities onto the SHAPE-constrained structure maps revealed single-stranded nucleotides were significantly more reactive than double-stranded nucleotides (**Fig S1I-L, S1U-X**). Together, this demonstrates that the predicted structures are supported by two orthogonal experimental probing methods.

We first examined the 5’ termini for the conserved hairpin containing the 12-nt TSS. While these hairpin motifs have been inferred from studies *in silico* and *in vitro*, their formation during infection has not been demonstrated (23, 42, 43). In all four structure maps, the TSS hairpin structures exactly match the previously predicted mRNA structures (**Fig 1C**) (23). Although reactivities were masked at the extreme 5’ primer binding sites, including the entire VP35 stem loop, downstream reactivities in the longer hairpins of VP40, VP30, and VP24 provided clear experimental support. Generally, predicted stems had low reactivities while loops and bulges had medium to high reactivities (**Fig 1C**). Despite the lack of reactivities in the VP35 TSS hairpin, the elevated downstream reactivities support that the rest of the 5’ UTR is mostly single-stranded, suggesting that VP35 TSS hairpin formation occurs independently of the downstream nucleotides (**Fig S2A**). Together, these findings provide the first evidence supporting 5’-end hairpin formation in EBOV mRNAs during infection.

Next, we compared the global structural content of EBOV mRNAs to well-characterized +ssRNA virus genomes (HCV, SARS-CoV-2, WNV, and DENV2) which function similarly as viral translation templates. We quantified the overall structural density of EBOV mRNAs by calculating the base pairing content (BPC), expressed as the fraction of nucleotides involved in base-pairing interactions within the SHAPE-constrained maps (44). Across all four transcripts, we consistently observed similar levels of base-pairing, averaging 48±2% (**Fig 1D**). While these levels indicate that the EBOV transcripts are folded, the global BPC is somewhat lower than that of +ssRNA viral genomes, which average 55±6% BPC, typically with more than half of all nucleotides in base-pairs. These data suggest that while nsNSV mRNAs are structured, they possess a slightly more flexible global architecture compared to +ssRNA viral genomes, potentially reflecting their specialized roles as templates for translation.

To identify the source of the global structural difference, we calculated BPC within individual mRNA domains: the 5’ UTR, CDS, and 3’ UTR. Remarkably, the BPC within all four EBOV mRNA coding regions averaged 55±4%, which is nearly identical to the structural density within +ssRNA viral genomes (avg: 55±6%) (**Fig 1E, center**). This suggests that high structural content within translation templates is a general feature across RNA viruses, likely coordinating efficient translation or providing transcript stability. In contrast, the UTRs account for the structural differences observed between EBOV and +ssRNA viruses. The average BPC within EBOV mRNA 5’ UTRs (40±9%) and 3’ UTRs (38±6%) was markedly lower than the +ssRNA virus counterparts, 63±8% and 51±9%, respectively (**Fig 1E**). The sole exception was VP40, which contained BPC comparable to +ssRNA virus 5’ UTRs; however, this is likely a result of having the shortest 5’ UTR (89 nt) and the 59-nt TSS hairpin accounting for two thirds of the 5’ UTR. Collectively, these data indicate that EBOV mRNAs prioritize high structural density in the CDS over UTRs, likely reflecting an adaptation for translation-only templates.

### EBOV mRNAs contain networks of structured RNA elements

Beyond global structural content, we also characterized the local architectural features of the EBOV mRNAs. To identify potential structure candidates for functional validation, we calculated the median SHAPE reactivity and Shannon Entropy in 51-nt sliding windows. Regions with low backbone flexibility (SHAPE reactivity) and low conformational heterogeneity (Shannon entropy), or “LowSS” have been established as hotspots for functional RNA elements across a variety of RNA viruses (9). We defined LowSS regions as stretches of >35 nucleotides where both local median SHAPE reactivity and Shannon entropy fall below the global median in both replicate datasets, consistent with the literature (**Fig 2A-D**) (9, 14, 45).

**Fig 2.**
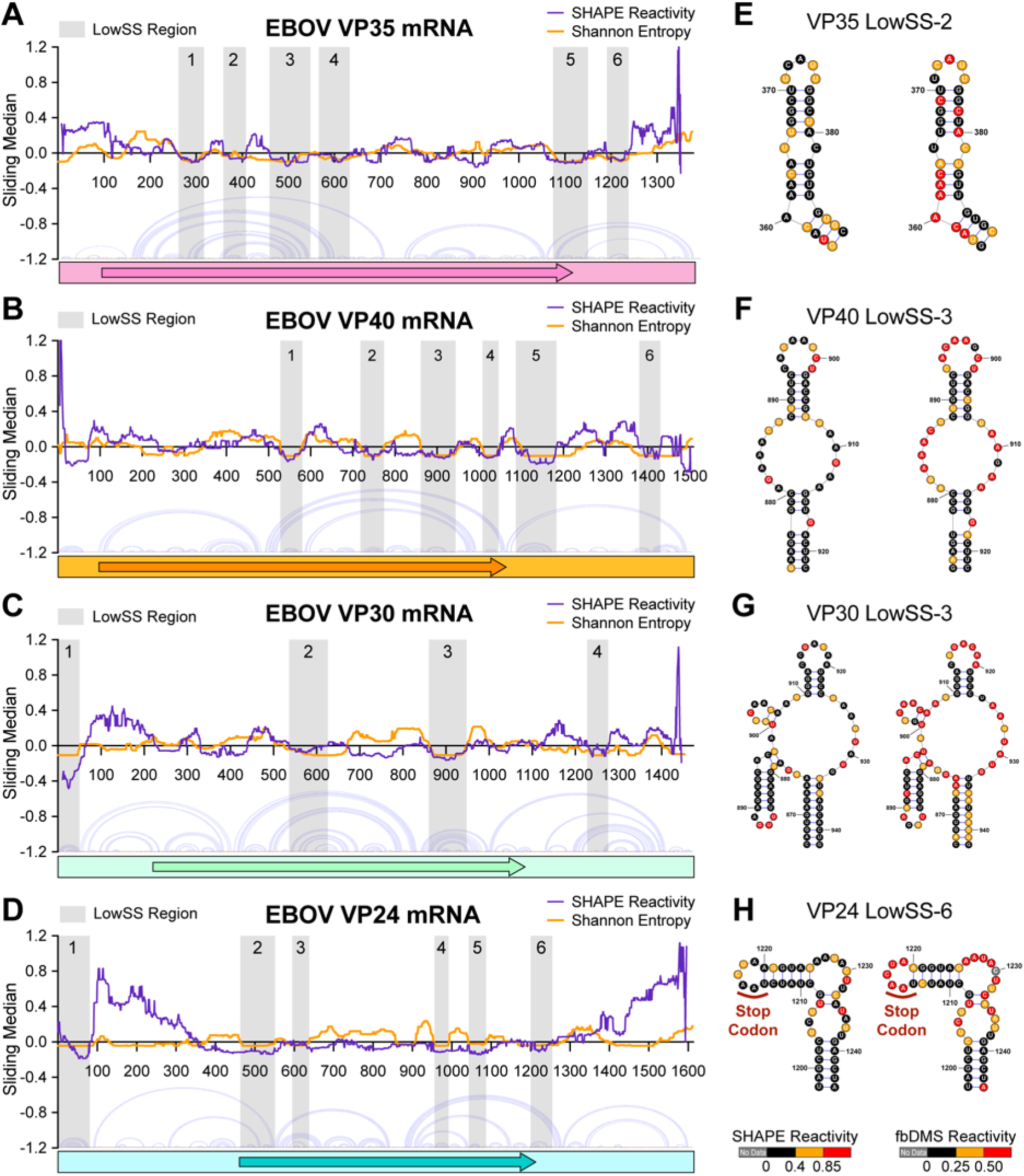
SHAPE-MaP and fbDMS-MaP reveal highly structured elements in EBOV mRNAs. (**A-D**) Analysis of local median SHAPE reactivity and Shannon entropy in (**A**) VP35 (1376 nt), (**B**) VP40 (1505 nt), (**C**) VP30 (1453 nt), and (**D**) VP24 (1612 nt) mRNAs, with nucleotide coordinates shown on the x-axis. Arc plots show SHAPE-predicted base pairs above mRNA schematics with shaded arrows denoting the CDS. Grey shading indicates stable structural regions where both SHAPE reactivity and Shannon entropy fall below the global median (LowSS). (**E-H**) Representative illustrations of SHAPE-constrained LowSS structures colored by (left) SHAPE 2A3 reactivity and (right) fbDMS reactivity.

The structural organization of VP35 and VP40 mRNAs revealed a distinctive clustering of LowSS elements. In VP35 and VP40 mRNAs, we identified six LowSS structures concentrated in two locations: the CDS and 3’ UTR (**Fig 2A-B, Fig S2A-B**). The CDS clusters contained four LowSS elements, while the 3’ UTR clusters contained two LowSS elements. The similar structural arrangement in the 3’ UTR with a LowSS element spanning or near the stop codon, followed by another LowSS element, suggests a regulatory array that potentially coordinates translation or transcription termination.

In contrast, LowSS elements were more distributed in the VP30 and VP24 mRNAs, across the 5’ UTR, CDS, and 3’ UTR. While the VP24 mRNA also had six LowSS elements, VP30 only had four LowSS elements, the fewest of the mRNAs (**Fig 2C-D, Fig S2C-D**). Both mRNAs had one LowSS structure in the 5’ UTR corresponding to the TSS hairpin, and one LowSS structure in the 3’ UTR. Interestingly, VP24 had two junction-spanning elements; LowSS-2 was located at the 5’ UTR-CDS junction, a feature unique to the VP24 transcript, and LowSS-6 that spanned the CDS-3’ UTR junction, like VP35 LowSS-5 (**Fig 2D**). These junction-spanning motifs suggest boundary-associated structures and regulatory mechanisms are maintained across transcripts despite the varying global distribution of LowSS elements.

Side-by-side comparisons of SHAPE and DMS reactivities mapped onto these LowSS structures demonstrate good agreement, indicating the formation of these elements is supported by both probing methods (**Fig 2E-H**). For some motifs like VP35 LowSS-2 and VP30 LowSS-3, DMS reactivities were slightly elevated in a subset of base-paired nucleotides (**Fig 2E, 2G**). This may reflect the elevated Shannon entropy within or flanking the LowSS region, indicating the presence of a few conformationally flexible nucleotides (**Fig 2A, 2C**). The higher median DMS mutation rates suggest that DMS may be more sensitive to transiently single-stranded states compared to SHAPE (**Fig S1A-D, S1M-P**). Overall, orthogonal cross-validation reinforces that these newly identified LowSS structures are stably formed during infection.

To our knowledge, this work provides the first comprehensive secondary structure maps of nsNSV and filovirus RNA architectures obtained during infection at BSL-4, and reveals new structural features that may be critical regulators of viral infection.

### EBOV mRNA 5’ stem loop structures are critical for infection

To evaluate the functional significance of the SHAPE-MaP structures, we used antisense oligonucleotides containing locked nucleic acids (LNAs) to disrupt the LowSS elements. LNAs are synthetic nucleotide analogs that increase the melting temperature (T_m_) of the oligonucleotide upon hybridization by 2-8°C per substituted nucleotide, which enhances binding affinity and facilitates disruption of endogenous base-pairing (46). LNA-mediated structure disruption is an established method for testing the functionality of SHAPE-predicted viral RNA motifs, as used for HCV, WNV, SARS-CoV-2, and influenza A virus (12, 14, 47–49).

To test whether the mRNA TSS structures regulate infection, we designed two types of LNAs targeting either the 12-nt TSS sequence (TSS-LNA) or the TSS hairpin structure (TSS-HP-LNA) (**Fig 3A**). The TSS is identical across VP35, VP40, GP, VP30, and VP24 mRNAs (5’-GA<u>U</u>GAAGAUUAA-3’) but it differs in NP and L mRNAs by one nucleotide (5’-GA<u>G</u>GAAGAUUAA-3’). To account for this variation, we synthesized 12-nt LNAs targeting both the VP-GP and NP-L TSS. To disrupt stem loop structures, longer TSS-HP-LNAs (20-30 nt) were designed using the SHAPE-constrained structures for each mRNA (**Fig 3A, right**). A scrambled LNA with no predicted complementarity to any EBOV RNAs was also included as a global negative control. All LNAs were synthesized as LNA/DNA mixmers to minimize cell toxicity, with three locked nucleotides at each end to enhance end stability, and no more than three consecutive DNA nucleotides to avoid RNase H degradation (50). Disruptions to viral infection were quantified by fluorescence microscopy using the rEBOV-ZsGreen-VP40 reporter virus (51).

**Fig 3.**
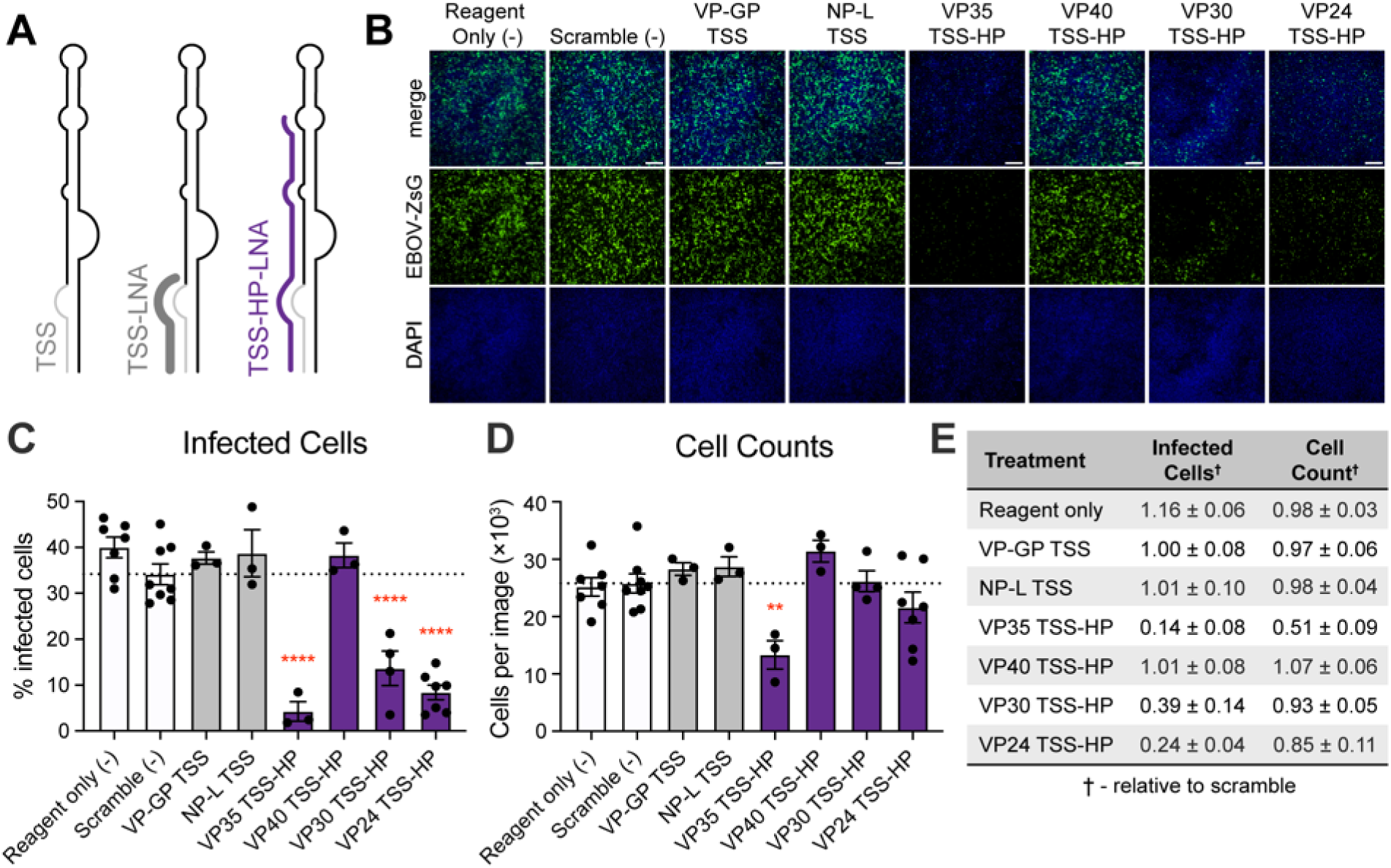
Reduction of EBOV infection upon LNA targeting of mRNA 5’ stem loops. (**A**) Schematics of the EBOV mRNA 5’ hairpin containing the TSS (left) and LNA disruption strategy targeting the 12-nt TSS (center) or the hairpin structure (TSS-HP right). (**B**) Representative fluorescence microscopy images of EBOV-infected Huh7 cells following LNA treatment. Cells were transfected with 400 nM LNA for 4-5 hours before infection (rEBOV-ZsGreen-VP40; MOI = 0.5). At 2 dpi, formalin-fixed cells were DAPI stained, and images were acquired at 4x magnification (scale bar = 500 µm). (**C-D**) Quantification of (**C**) infected cells and (**D**) total cell counts after LNA treatment using QuPath. Bars represent global negative controls (white), TSS-LNAs (grey), and structure-disrupting LNAs (violet). Data points indicate mean ± SEM of (n≥3) biological replicates (performed in technical duplicate, ≥4 images per sample). Statistical significance relative to scramble LNA by ordinary one-way ANOVA: ** p < 0.01; **** p < 0.0001. (**E**) Average change (± SEM) in infected cells or cell counts following LNA treatment, relative to the scrambled LNA. VP-GP TSS denotes the 12-nt TSS shared by VP35, VP40, GP, VP30, and VP24 mRNAs; NP-L TSS denotes the 12-nt TSS in NP and L mRNAs.

First, we tested whether targeting the TSS in EBOV mRNAs affected infection. Both 12-nt TSS-LNAs (VP-GP and NP-L) had little to no effect on infection or cell counts (**Fig 3B-E**). Several factors could have contributed to the lack of inhibition: the 12-nt LNAs might have been too short to anneal, they could have broad targeting of shared sequences, or the TSS might only have regulatory function in the viral genome.

In contrast, infection was significantly reduced by the longer LNAs targeting the TSS hairpin structures in VP35 (85%), VP30 (61%), and VP24 (76%), though targeting VP35 also caused a 49% reduction in cell numbers (**Fig 3C-E**). Surprisingly, the VP40 TSS-HP LNA had no effect on infection, potentially due to bound proteins protecting the stem from disruption, or other environmental factors preventing LNA hybridization (**Fig 3B-C**). Together, these results demonstrate that the conserved TSS hairpin structures in EBOV mRNAs are critical for infection.

### Disrupting mRNA secondary structures reduces EBOV infection

Next, we examined whether the newly identified LowSS elements also regulate viral infection. We designed LNAs to outcompete one side of a predicted stem to maximize structure disruption while avoiding self-hybridization. For negative controls, LNAs were also designed to target ssRNA in the same region (5’ UTR, CDS, or 3’ UTR), prioritizing nucleotides with high SHAPE and DMS reactivities where targeting was unlikely to affect infection. This systematic approach enabled us to identify one to three regulatory structures per mRNA that significantly reduced infection with minimal LNA cytotoxicity (**Fig S3A-D**).

In the VP35 mRNA, we identified three functional LowSS elements, LowSS-2, LowSS-5, and LowSS-6. Disrupting LowSS-5 and LowSS-6 in the 3’ UTR significantly reduced infection by 57% and 51%, respectively (**Fig 4A**). These structures potentially play interconnected roles given their shared location and similar LNA phenotypes. In contrast, disrupting LowSS-2 in the CDS yielded an intermediate reduction of 41% (**Fig 4A, Fig S4E**). Since LowSS-2 contains flexible nucleotides with elevated Shannon entropy (nts 382-385), LNA targeting may only disrupt one functional conformation while alternative structure(s) remain intact and preserve viral function (**Fig 2A**). A similar phenomenon occurs with the SARS-CoV-2 ribosomal frameshifting pseudoknot, where LNA targeting slightly reduced infection by 17-18% (12). Since ssRNA negative controls did not disrupt infection, this indicates the observed reductions reflect structure-specific targeting (**Fig S4A, S4E**). Overall, this demonstrates that EBOV mRNAs harbor functional secondary structures across coding and untranslated regions, likely reflecting diverse regulatory mechanisms across the viral life cycle.

**Fig 4.**
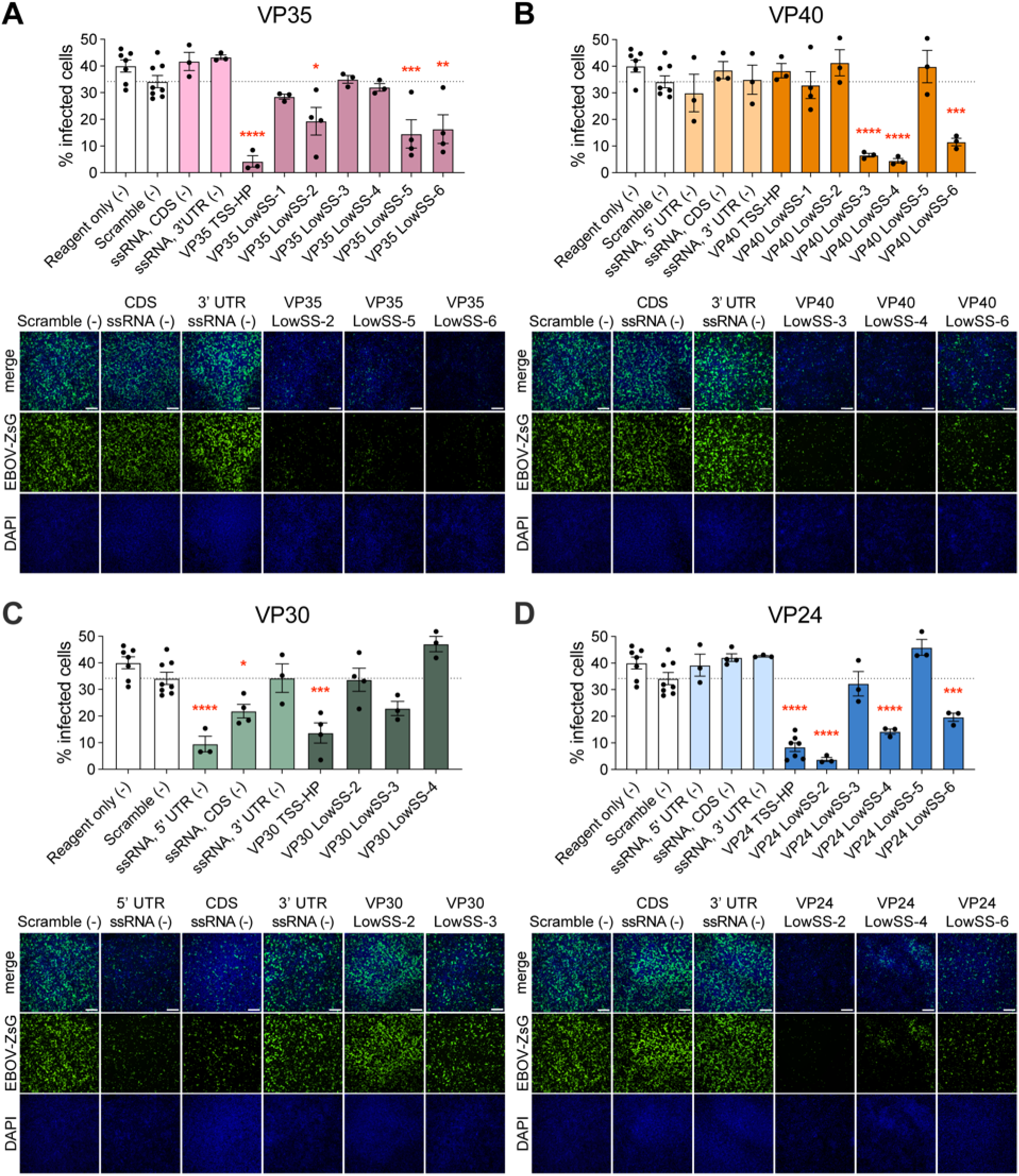
LNA targeting of EBOV LowSS structures reduces EBOV infection. (**A-D**, top) Quantification of infected cells upon LNA treatment of (**A**) VP35, (**B**) VP40, (**C**) VP30, and (**D**) VP24 mRNAs. Data points represent the mean infection rate ± SEM from (n≥3) biological replicates (performed in technical duplicate; ≥4 images per replicate). Bars represent global negative controls (white), LNAs targeting ssRNA (lighter colors), and structure-disrupting LNAs (darker colors). Reagent-only and scramble negative controls are reproduced from Fig 3, as experiments were performed simultaneously with the same set of controls. TSS hairpin (TSS-HP) LNA data are reproduced from Fig 3 to compare previously predicted structures and newly identified LowSS structures. Dotted lines indicate mean infection rate after scramble LNA treatment (33.4%). Statistical significance relative to scramble determined by ordinary one-way ANOVA: * p < 0.05; ** p < 0.01; *** p < 0.001; **** p < 0.0001. (**A-D**, bottom) Representative fluorescence microscopy images of EBOV-infected Huh7 cells following LNA targeting. Cells were transfected with 400 nM LNA for 4-5 hours before infection (EBOV-ZsGreen-VP40; MOI = 0.5), fixed at 2 dpi with formalin and DAPI-stained (4x magnification, scale bar = 500 µm).

In the VP40 mRNA, we also found three LowSS motifs in the CDS and 3’ UTR that were critical for infection. Specifically, LNA disruption of LowSS-3, LowSS-4, and LowSS-6 significantly reduced infection by 80%, 87%, and 64%, respectively, with minimal off-target activity (**Fig 4B, Fig S4B**). Both VP40 LowSS-3 and LowSS-4 are small single stem-loop structures in the CDS that yielded the strongest reductions in infection in this study despite their small size and simple architecture (**Fig S2B, S4F**). These data demonstrate that simple CDS stem loop structures in EBOV mRNAs can function as critical regulators of viral infection.

For VP24, disrupting LowSS structures at three distinct loci reduced infection across a range of 88% (LowSS-2, 5’ UTR-CDS junction), 56% (LowSS-4, CDS), and 39% (LowSS-6, CDS-3’ UTR junction) (**Fig 4D; Fig S4H**). Though VP24 LowSS-2 disruption reduced infection to 12%, this LNA also caused off-target reductions in cell viability by 55% (**Fig S4D**). Nonetheless, these findings suggest the regulatory elements in the VP24 mRNA are strategically organized at key transcript junctions, possibly regulating translation initiation or termination.

Unlike the previous mRNAs, targeting the VP30 LowSS structures had minimal impact on infection. Only LowSS-1 (TSS hairpin) disruption significantly reduced infection (**Fig 4C, Fig S4G**). Targeting the LowSS-3 structure led to a modest decrease in infection of 31%, possibly due to nearby regions with increased Shannon entropy (**Fig 2C, 4C**). Unexpectedly, targeting ssRNA in the 5’ UTR (nts 168-189) and CDS (823–844) significantly reduced infection by 69% and 35%, respectively, suggesting VP30 contains unannotated ssRNA regulatory elements (**Fig 4C, Fig S4G**). These LNA treatments did not affect cell viability, indicating disruption is EBOV-specific (**Fig S4C**). Together, these findings show that the VP30 mRNA is unusual in that it contains more flexible regulatory elements than the other mRNAs, which may adopt multiple conformations that tune expression levels of the VP30 transcription factor.

Overall, these results demonstrate that EBOV mRNAs contain a diverse repertoire of static, dynamic, and single-stranded architectures across key junctions to form elaborate regulatory networks that are critical for EBOV infection.

### Evolutionary support for RNA structures across mammalian filoviruses

After identifying functional RNA structure candidates by LNA targeting, we asked whether these elements exhibit evolutionary conservation across the filovirus family. Synonymous mutation rates (SMR) reveal evolutionary pressure on viral RNA structures, since mutations disrupting critical base-pairing interactions are deleterious even if they do not alter the protein sequence (11–13). While new piscine and reptilian filoviruses are being discovered, mammalian filoviruses remain the primary threats to human health (52, 53). Therefore, we analyzed SMR using VP35, VP40, VP30, and VP24 coding sequences from only mammalian filoviruses (**Fig 5A**). We observed significantly lower SMR in base-paired nucleotides for VP35, VP30, and VP24, indicating that these coding sequences are under evolutionary pressure to maintain their structure content across mammalian filoviruses (**Fig 5B**).

**Fig 5.**
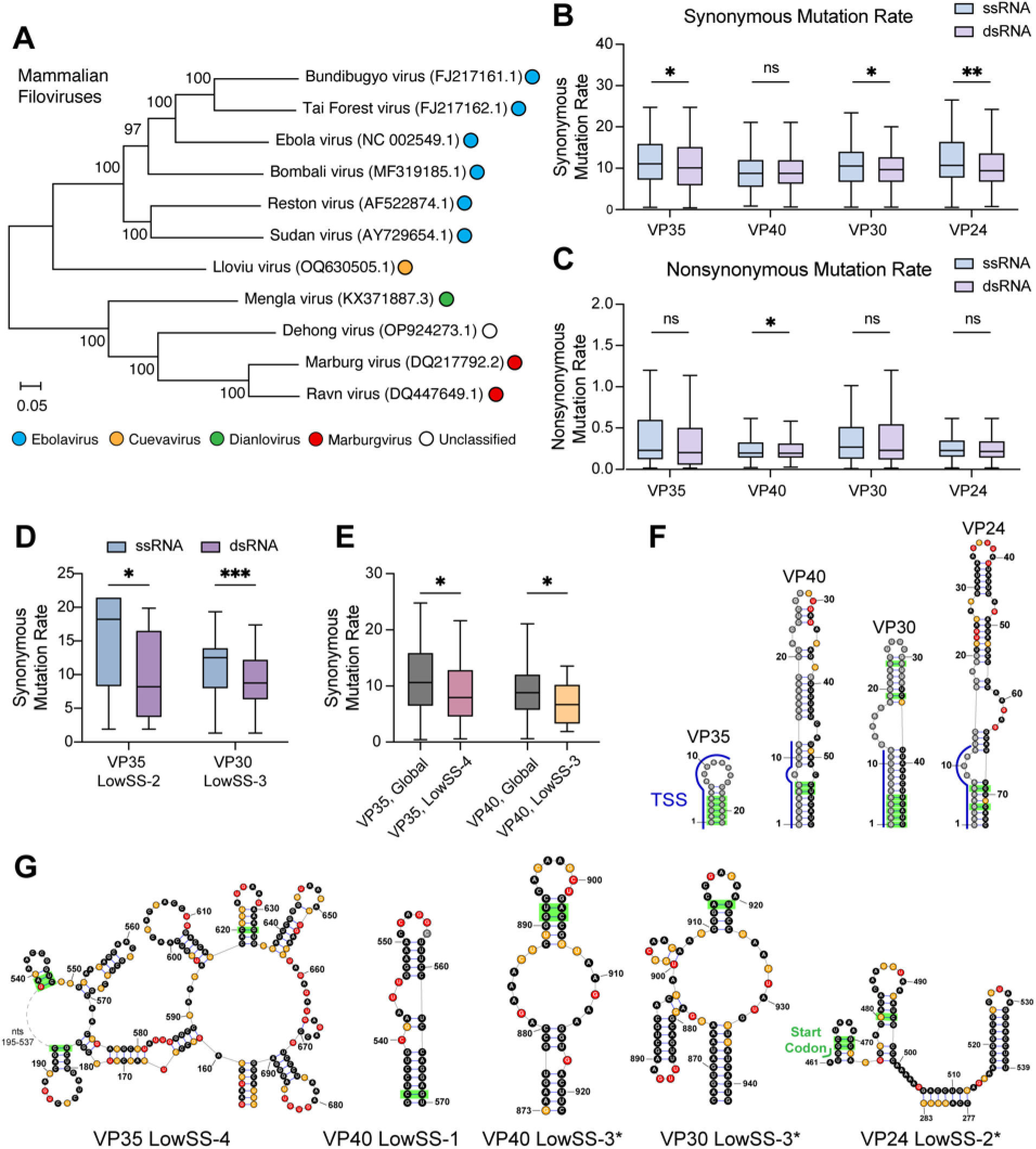
Synonymous mutation rate and covariation provide evolutionary support for functional EBOV RNA structures in mammalian filoviruses. (**A**) Phylogenetic tree computed from a multiple sequence alignment (MSA) of all 11 mammalian filoviruses genomes using the general time reversible model (4500 bootstrap replicates). Node labels indicate bootstrap support values and scale bar indicates evolutionary distance in nucleotide substitutions per site. Member viruses are color-coded by genus. (**B-C**) (**B**) Synonymous and (**C**) Nonsynonymous mutation rates calculated from an MSA of mammalian filovirus coding sequences, mapped onto single– and double-stranded nucleotides of the SHAPE structure model. Data are shown in Tukey style with outliers omitted (boxes: interquartile range (IQR); whiskers: 1.5 x IQR; horizontal line: median). (**D-E**) Synonymous mutation rates calculated from mammalian filovirus CDS alignments (**D**) mapped onto single– and double-stranded nucleotides of LowSS regions, or (**E**) in whole LowSS regions compared to the global CDS synonymous mutation rate, plotted as in (**B-C**). (**F-G**) EBOV (**F**) TSS hairpin and (**G**) LowSS structures containing significantly covarying base pairs (green) analyzed by R-scape (--RAFSp, Window: 200, Slide: 100, E < 0.05) for a mammalian filovirus mRNA alignment. LowSS structures with LNA phenotypes of >30% reduction in infection are indicated by an asterisk (*). Statistical significance: ns, not significant; * p < 0.05; ** p < 0.01; *** p < 0.001; by two-tailed unpaired Student’s t-test (**B-D**) or ordinary one-way ANOVA (**E**).

In contrast, VP40 exhibits the opposite evolutionary profile. No significant differences were observed between SMR in ssRNA and dsRNA (**Fig 5B**). Interestingly, nonsynonymous mutation rates appear to be slightly suppressed in base-paired regions (**Fig 5B-C**). This suggests the VP40 CDS has different evolutionary constraints than the other mRNAs. This observation aligns with the evolution of the VP40 matrix protein within mammalian filoviruses over time. VP40 drives virion assembly and budding in filoviruses, but Marburg and Měnglà virus VP40 proteins additionally inhibit IFN signaling, a function absent in EBOV and Lloviu virus VP40 (54–56). VP40 diverges further across the filovirus family, with homologs absent from the newly discovered piscine filoviruses (7).

To refine our analysis, we then analyzed SMR within the individual EBOV LowSS elements to identify evolutionary support for these structures across mammalian filoviruses. Two structures, VP35 LowSS-2 and VP30 LowSS-3, exhibited significantly lower SMR in dsRNA compared to ssRNA (**Fig 5D**). Interestingly, LNA disruption of both structures yielded intermediate infection reductions, and VP30 LowSS-3 contains a substructure (nts 878-898) with similarity to VP35 LowSS-2 (**Fig 2E, 2G**). The convergence of structural homology, evolutionary constraint, and LNA disruption phenotype suggests that these motifs represent a conserved class of regulatory structures in mammalian filoviruses.

Regional suppression of synonymous mutations, irrespective of base-pairing status, can also reveal additional evolutionary constraints on LowSS structures. We identified two structures, VP35 LowSS-4 and VP40 LowSS-3, in regions with significantly suppressed SMR compared to the whole CDS (**Fig 5E**). This selection indicates regulation occurs at both sequence and structure levels, like the TSS hairpin in the genomic 3’ leader which initiates EBOV mRNA transcription (22). The sequence and structure constraints at these regions indicate these particular RNA elements may be amenable to sequence-based attenuated viral vaccines or antisense disruption therapeutic strategies.

To obtain direct evidence of base-pair conservation, we performed covariation analysis using mRNA sequence alignments from all eleven mammalian filoviruses. We identified significantly covarying base pairs in the four TSS hairpins and in five LowSS structures (**Fig 5F-G**). Two of the elements with LNA disruption phenotypes, VP40 LowSS-3 and VP30 LowSS-3, had both SMR and covariation support, providing strong evidence for conservation across mammalian filoviruses (**Fig 5D-E**). In contrast, VP35 LowSS-4 and VP40 LowSS-1 contained covarying base pairs, but lacked LNA disruption phenotypes (**Fig 5G**). These conserved structures may still be functional, but inaccessible to LNA targeting, like the VP40 TSS hairpin. Altogether, the convergence of evolutionary support by SMR and covariation suggests that these seemingly simple stem loop structures form the functional cores of important EBOV regulatory elements.

We also found significantly covarying base pairs outside of LowSS structures. Covariation was observed in the stem adjacent to the VP24 LowSS-4 structure that displayed an LNA disruption phenotype (**Fig S5A**). This covariation near a LowSS structure suggests the formation of local LowSS motifs that are embedded within larger, conserved parental structures. We also found three significantly covarying base pairs in a VP24 CDS structure (nts 760-830) that displays low SHAPE reactivity and high Shannon entropy (**Fig 2D, Fig S5B**). This type of signature, where chemical reactivity is low, but entropy is high, strongly suggests formation of a conserved regulatory switch capable of adopting multiple conformations.

Altogether, this study reveals that mammalian filoviruses contain a complex repertoire of static and dynamic regulatory elements that function through both primary sequence and secondary structure. These new functional RNA elements not only advance our understanding of filovirus infection, but also serve as new high-priority targets for pan-filoviral therapeutics.

## Discussion

In this study, we experimentally mapped the secondary structures of four EBOV mRNAs in infected cells and demonstrated that LNA targeting of these structures significantly reduces infection. This work addresses the limitations of *in silico* and *in vitro* systems by integrating standard chemical probing into BSL-4 settings and provides a foundation for mechanistic studies of filovirus replication. More broadly, we extend in-cell RNA secondary structure analysis to nsNSVs, a viral order with vastly different genomic architectures and life cycles than previously characterized viruses.

The distinct replication strategies of nsNSVs and +ssRNA viruses are likely reflected in the global architectures of their positive-sense transcripts. We find that EBOV mRNAs have highly base-paired CDS regions and less structured UTRs, while +ssRNA viruses are highly structured in both the CDS and UTRs (**Fig 1E**). The high propensity for structure formation throughout +ssRNA genomes is likely optimized to coordinate all viral processes, such as translation, replication, and packaging, from one RNA molecule (57, 58). nsNSVs like EBOV separate the processes of genome replication/transcription from translation by using mRNAs. Therefore, the lower UTR structure content likely reflects an optimization in mRNAs from nsNSVs for exclusively recruiting host translation machinery without having to contend with replication machinery. This indicates that nsNSVs may have evolved different RNA structure contents between translation and replication templates. This further suggests that while UTR complexity reflects the demands of specific viral replication strategies, high CDS structural content is a general feature of translated viral RNAs. However, confirming this multi-layered organization will require further in-cell structural mapping of the remaining mRNAs (NP, GP, L) and the negative-sense genome.

The broader architectural differences between positive– and negative-sense RNA viruses are also reflected in the local structures. In this study, we validated previously predicted EBOV mRNA TSS hairpin structures while revealing new regulatory elements in the CDS, 3’ UTR, and CDS-UTR junctions. These data indicate that functional structures are more pervasive in EBOV mRNAs than previously recognized. Notably, most of the newly identified functional elements are simple stem loops or stem loop arrays, rather than complex multi-loop architectures. These simple LowSS structures also tended to have AU-rich loop elements, mirroring previously identified regulatory elements, like the TSS hairpins, HSPA8 hairpin, and GP editing site structures (23, 27, 28). Viral genomes and antigenomes in nsNSVs are also tightly encapsidated by nucleoproteins, a constraint that may limit the complexity of RNA secondary structures in both the negative– and positive-sense RNAs. Likewise, LowSS structures in influenza A virus, a segmented negative-sense RNA virus, tend to adopt simple hairpin architectures (16). This pattern suggests negative-sense RNA viruses may contain simpler regulatory elements compared to the more complex multi-loop or pseudoknot structures commonly found in +ssRNA viruses, potentially reflecting distinct replication mechanisms between these viral phyla (58).

Little is known about the regulatory roles of RNA secondary structures during nsNSV replication, transcription, and translation, nor how these functions are divided between the genome, antigenome, and mRNAs. The locations of the EBOV mRNA secondary structures may hint at their roles; for example, 3’ UTR structures may be involved in terminating transcription, while the CDS structures regulate processes related to translation, though such mechanisms have yet to be experimentally evaluated. The clustering of functional structures at the 3’ region of VP35 and VP40 mRNAs suggests a regulatory array that forms at the boundaries of high-abundance transcripts. VP35 LowSS-5/6 and VP40 LowSS-6 are arrays of 3’ UTR stem loops that produced similar functional phenotypes, reducing infection by 50-65% upon LNA disruption (**Fig 4A-B**). Filovirus UTRs have been hypothesized to contain integral regulators of transcription and transcript abundance through *cis*-acting sequence and structure elements (59). These LowSS structures may modulate RdRp processivity in highly transcribed genes to facilitate polyadenylation and transcription termination, like prokaryotic Rho-independent systems (59–61). Together, these arrays likely constitute a distinct class of elements involved in transcript processing, a critical regulatory handoff for nsNSV gene expression that currently remains mechanistically unresolved.

We also identified functional structures at CDS-UTR boundaries, such as VP24 LowSS-2 at the 5’ UTR-CDS boundary, and VP35 LowSS-5 and VP24 LowSS-6 at the CDS-3’ UTR boundary. The junction-spanning structures may modulate ribosomes in a locus-dependent manner, whereby 5’ UTR-CDS structures could enhance translation, while CDS-3’ UTR structures might terminate translation. Notably, these types of structures were only identified in VP35 and VP24 mRNAs, both of which encode immunomodulatory nucleocapsid proteins, suggesting that translation regulation at CDS boundaries may control the critical expression of immunosuppressive viral factors (8). The presence of functional LowSS elements at the CDS-UTR boundaries is also reminiscent of the regulatory architectures in +ssRNA viruses, where boundary structures induce ribosomal pausing (13, 14, 62, 63). For example, pseudoknot structures in the genomes of SARS-CoV-2 (nsp10-nsp11) and HCV (NS2-NS3) act as physical barriers that coordinate frameshifting or downregulation of translation (64, 65). This suggests that, like +ssRNA viruses, EBOV coding transcripts may also use regulatory elements at CDS boundaries to control ribosomal progression.

In many cases, LowSS structures were identified in the middle of the CDS, for which LNA treatment reduced infection, and both SMR and covariation suggested conservation across mammalian filoviruses. The agreement between functional and evolutionary data in VP35 LowSS-2, VP40 LowSS-3, and VP30 LowSS-3 indicates that these elements modulate critical viral processes and represent viable candidates for pan-filoviral therapeutics. In +ssRNA viruses, CDS structures are known to regulate translation and inhibit host immune responses (64–66), and elements in the EBOV CDS regions may have analogous regulatory roles. While the regulatory landscape of nsNSV RNA structures is still largely uncharacterized, direct mechanistic characterization of RNA elements will be required to elucidate their functions.

Beyond static LowSS elements, this study reveals that EBOV mRNAs may harbor conformationally flexible structures that contribute to viral fitness. While LowSS is an established metric for identifying static candidate functional structures, this work supports the importance of identifying RNA motifs that behave as dynamic structural ensembles (33, 67, 68). Structures like VP35 LowSS-2 and VP30 LowSS-3 have high Shannon entropy nucleotides and displayed intermediate effects upon LNA targeting, but there is evolutionary support for their importance by suppressed SMR and covariation. Traditional LowSS analyses may overlook motifs that contain more than one stable conformation, and which might function as switches. The additional motif in the VP24 CDS is one such candidate (**Fig S5B**). Although conformational flexibility may yield intermediate antisense targeting effects, alternative drug modalities like small molecules may be highly effective against such flexible, conserved RNA elements. Together, the conservation of switch-like structures across mammalian filoviruses suggests that conformational flexibility may be essential for coordinating viral gene expression, potentially through uncharacterized higher-order RNA-RNA interactions or ribonucleoprotein complexes.

Taken together, our findings establish the location and importance of static and dynamic secondary structural elements within EBOV mRNAs. We show that filovirus mRNAs possess networks of simple but critical regulatory structures, reinforcing that positive-sense viral transcripts efficiently encode functionality within their primary sequences and secondary structures. This analysis of EBOV RNA elements deepens our understanding of filovirus infection and expands the pool of potential targets for pan-filoviral therapeutics.

## MATERIALS AND METHODS

### Biosafety statement

All work with Ebola virus (EBOV) was performed in the biosafety level 4 (BSL-4) facility of Boston University’s National Emerging Infectious Diseases Laboratories (NEIDL) following approved standard operating procedures in compliance with local and federal regulations pertaining to the handling of BSL-4 pathogens and Select Agents.

### Cell Lines

African green monkey kidney cells (Vero E6; ATCC: CRL-1586) and Huh7 cells (kindly provided by Apath L.L.C.) were maintained in Dulbecco’s modified Eagle medium (DMEM; Thermo Fisher Scientific) supplemented with 200 mM L-glutamine (Thermo Fisher Scientific), 100 μg/mL Primocin (Invivogen), and 10% fetal bovine serum (FBS; R&D Systems). Cells were cultured at 37 °C and 5% CO_2_.

### Viruses

EBOV (isolate Mayinga, RefSeq: NC_002549.1) was generously provided by H. Feldmann, Rocky Mountain Laboratories, NIAID, NIH. EBOV-ZsGreen (GenBank: OL956940.1) was generated as previously described (69). Virus stocks were propagated in Vero E6 cells in DMEM supplemented with 200 mM L-glutamine, 100 μg/mL Primocin, and 2% FBS and purified by ultracentrifugation through a 20% sucrose cushion as previously described (70). Virus titers were subsequently determined in Vero E6 cells by tissue culture infectious dose 50 (TCID_50_) assay using the Spearman and Kärber algorithm (71).

### EBOV infection of cells for chemical probing

Huh7 cells were seeded at a density of 5×10^6^ cells into T175 tissue culture flasks. The next day, cells were infected with EBOV at a multiplicity of infection (MOI) of 5 by adding 20 mL of DMEM supplemented with 200 mM L-glutamine, 100 μg/mL Primocin, and 2% FBS containing the appropriate volume of EBOV stock solution. Cells were incubated for 1 d at 37°C and 5% CO_2_ until harvest for chemical probing.

### SHAPE chemical probing

Within 2 h of probing, a 1 M stock solution of 2A3 was prepared by resuspending 5 mg of 2A3 (kindly provided by New England Discovery Partners) in 26.5 µL of anhydrous DMSO (Sigma-Aldrich) per biological replicate (37). At 1 day post-infection (dpi), cell supernatant was removed from EBOV-infected flasks and cells were washed with 10 mL of room temperature PBS. Cells were dislodged from the flasks using large cell scrapers, collected in a 15 mL conical tube, and centrifuged at 500 × g for 5 min at 4°C. Each cell pellet was resuspended in 250 µL of PBS, and the suspensions from two flasks were pooled for biological replicates and then separated again into two 250 µL suspensions for probing. Each cell suspension was treated with either 25 µL of DMSO or 2A3 in DMSO (final concentration = 100 mM) and mixed by repeatedly pipetting. Samples were incubated for 15 min at 37°C. To facilitate probe diffusion, cells were mixed again halfway through the incubation by inverting tubes several times. Following incubation, samples were spun down at 500 × g for 3 min. The supernatant was removed, and cell pellets were resuspended in 2 mL TRIzol (Invitrogen) following approved inactivation SOPs, removed from the BSL-4 laboratory, and stored at –80°C until RNA purification.

### fbDMS chemical probing

Within 2 h of probing, a stock solution of 1.7 M dimethyl sulfate (DMS) was prepared by mixing 26.8 µL of 100% DMS (Sigma-Aldrich) into 140 µL of 200 proof ethanol (Fisher Scientific) per biological replicate. DMS treatment of EBOV-infected cells was performed at 1 dpi as described above for SHAPE chemical probing with the following differences: Each cell pellet was resuspended in 1 mL of room temperature 1 M bicine buffer (pH 8.3), pooled for biological replicates, then divided again into two 1 mL suspensions for probing. Each cell suspension was treated with either 100 µL of ethanol or DMS in ethanol (final concentration = 170 mM) and mixed by pipetting repeatedly. Samples were incubated for 6 min at 37°C. To facilitate probe diffusion, cells were mixed again halfway through the incubation by inverting tubes several times. Following incubation, 3.3 mL of TRIzol LS (Invitrogen) was added to each sample following approved inactivation SOPs. The samples were removed from the BSL-4 laboratory and stored at –80°C until RNA purification.

### RNA purification

RNA was extracted by mixing 24:1 chloroform:isoamyl alcohol (Millipore Sigma) at a ratio of 1:5 volumes of chloroform:isoamyl alcohol to sample for SHAPE samples in TRIzol or 1:3.75 volumes of chloroform:isoamyl alcohol to sample for fbDMS samples in TRIzol LS and incubating at room temperature for 5 min. Samples were centrifuged at 12,000 × g for 15 min at 4°C, and the aqueous phase was collected in a new tube. RNAs were precipitated by adding 3 volumes of 100% ethanol and incubated overnight at –20°C. The next day, RNAs were centrifuged at 12,000 × g for 1 h at 4°C. After centrifugation, the supernatant was aspirated, and the RNA pellet was resuspended in 100 µL of ME buffer (10 mM MOPS pH 6.5, 0.1 mM EDTA pH 8.5) and further purified using the QIAGEN RNeasy Mini kit (QIAGEN) according to the manufacturer’s protocol. RNA was eluted in 50 µL of ME buffer, and the RNA concentration was quantified using a Nanodrop 2000.

### Reverse transcription and PCR amplification of cDNAs

By utilizing the ultra-processive reverse transcriptase MarathonRT, RT-PCR primers were designed to capture the entire mRNA in a single PCR amplicon (**Table S1**) (39, 40). MarathonRT reverse transcriptase was expressed and purified as previously described (39). For each of the mRNAs, 500 ng of total cellular RNA was annealed with 1 µL of 1 µM gene-specific reverse transcription primer in a total reaction volume of 7.5 µL at 68°C for 5 min, then cooled to 12°C for 5 min on a Bio-Rad Thermocycler (**Table S1**). Then, reverse transcription was performed using MarathonRT in MaP buffer (50 mM Tris-HCl pH 7.5, 200 mM KCl, 5 mM DTT, 0.5 mM dNTPs, 1 mM MnCl_2,_ 20% glycerol, and 10 U of MarathonRT) at a final reaction volume of 20 µL for 3 h at 42°C. Reverse transcription was directly followed by MarathonRT inactivation at 70°C for 5 min. RNAs were degraded by adding 1.5 µL of a 1:1:1 mixture of RNase H (New England Biolabs, NEB), RNase A (NEB), and RNase T1 (Thermo Fisher Scientific) and incubating for 30 min at 37°C. The cDNAs were purified using AMPure XP beads (Beckman Coulter) using a 1.8x beads-to-sample ratio according to the manufacturer’s protocol. cDNAs were eluted in 13 µL of nuclease-free water and 5 µL cDNA was PCR-amplified with Q5 High-Fidelity DNA polymerase (NEB) using gene-specific primers according to the manufacturer protocol (**Table S1**). PCR amplicons were purified with AMPure XP beads (Beckman Coulter) using a 1.8x beads-to-sample ratio according to the manufacturer’s protocol. Cleaned amplicons were visualized on a 1% agarose gel to confirm production of the desired products prior to library preparation.

### MaP library preparation and sequencing

PCR amplicon concentrations were determined using the Qubit dsDNA HS assay (Thermo Fisher Scientific) and then diluted to 0.2 ng/µL. Sequencing libraries were prepared with the Nextera XT DNA Library Preparation kit (Illumina) according to the manufacturer’s protocol with the following changes: reactions were scaled down to 1/5^th^ the recommended volumes, samples were tagmented for 10 min at 55°C instead of 5 min, and Nextera PCR-amplified libraries were purified with AMPure beads using a 1.8x beads-to-sample ratio according to the manufacturer’s protocol, with an elution volume of 27 µL of nuclease-free water. Sample concentrations and the average library lengths were determined using Qubit HS dsDNA assay (Thermo Fisher Scientific) and a 2100 BioAnalyzer using the High Sensitivity DNA kit (Agilent). Libraries were diluted to 2 nM for pooling and loaded at a final concentration of 750 pM for sequencing on a NextSeq 2000 platform using the 300-cycle P1 kit (Illumina) according to the manufacturer’s protocol.

### Chemical probing data analysis and structure prediction

All SHAPE and DMS data were analyzed using ShapeMapper2 (v. 2.2) (41). Sequencing reads were aligned to the EBOV Mayinga mRNA reference sequences with the primer regions masked: VP35 (GenBank: NC_002549.1:3032-4407), VP40 (NC_002549.1:4390-5894), VP30 (NC_002549.1:8288-9740), and VP24 (NC_002549.1:9885-11496). For fbDMS data analysis, ShapeMapper2 was run with the minimum reactivity normalization factor threshold set to 0.0 (Shapemapper2/internals/bin/normalize_profiles.py, line 159). Default ShapeMapper quality control settings were used as a benchmark, and Pearson correlations for reactivities from independent biological replicates were calculated in GraphPad Prism as an additional quality control. SHAPE-constrained structure predictions were generated using SuperFold 1.2 using the default parameters (38). All secondary structure maps were visualized in StructureEditor (72).

### Base Pairing Content (BPC) calculations

To quantify the global structural density of EBOV mRNAs, we calculated base-pairing content (BPC) across the CDS and UTRs. BPC (± standard deviation, SD) was defined as the fraction of nucleotides involved in base pairs within the SHAPE-constrained structure for either the full-length mRNA, or individual regions (5’ UTR, CDS, or 3’ UTR) (44). For comparative analysis, BPCs in +ssRNA viruses were calculated using structures obtained from published in-cell SHAPE-MaP datasets: Dengue virus, serotype 2 (DENV2) (11); SARS-CoV-2 (12); Hepatitis C virus subgenomic replicon (HCV SGR) (13); West Nile Virus (WNV) (14).

### Identification of Low Shannon, Low SHAPE (LowSS) regions

To identify functional RNA structure candidates, local median Shannon entropies and SHAPE reactivities were calculated in 51 nt centered sliding windows for two independent SHAPE-MaP datasets. SHAPE reactivity and Shannon entropy values were obtained as outputs from ShapeMapper and SuperFold, respectively. The global median SHAPE reactivity or Shannon entropy value was subtracted from all local median values to normalize all data about the global median during visualization. Regions with local SHAPE reactivity and Shannon entropy below the global median for stretches longer than 35 nt and appeared in both independent replicates were considered “LowSS” (**Table S2**). Regions with short interruptions where reactivity or entropy rose above the global median for stretches of fewer than 35 nt were not disqualified.

### LNA design, synthesis and purification

Antisense oligonucleotides containing locked nucleic acids (LNAs) were designed to be between 20-40 nt long and target LowSS structures or single-stranded RNA regions in the mRNAs (**Table S3**). LNA mixmers contain three consecutive locked nucleotides at the 5’ and 3’ termini unlocked nucleotides, with a mix of locked and unlocked nucleotides in the center. A maximum of allowance of three consecutive unlocked nucleotides was used. A/T nucleotides were prioritized for locking, and all LNAs were designed to have similar thermodynamic properties: DNA:RNA duplex Tm ≥80°C, GC content >30%, and LNA content between 50-80%. LNAs were synthesized on a MerMade 12 synthesizer (BioAutomation) using DNA and LNA phosphoramidites (TX BIO) on Glen UnySupport™ 1000 controlled pore glass solid supports (Glen Research). Synthesis was performed according to the instrument manufacturer’s recommendation with a modified increased oxidation time of 3 min in the synthesis cycle. Base deprotection and oligonucleotide removal from CPG solid support were performed in a 1:1 mixture of 30% ammonium hydroxide (Fisher Scientific) and 40% aqueous methylamine (Millipore Sigma) at 65°C for 2 h. Deprotected oligo solutions were subsequently dried using a Savant™ Speedvac™ (Fisher Scientific) and desalted on a Glen Gel-Pak™ 1.0 Desalting Column (Glen Research). All final oligo concentrations were determined by Nanodrop 2000. Oligonucleotides were externally validated by Novatia LLC by mass spectrometry with electrospray ionization on a LTQ XL™ Linear Ion Trap Mass Spectrometer (Thermofisher), and for purity on a Vanquish UHPLC (Thermofisher).

### LNA structure disruption in EBOV-ZsGreen

Prior to transfection, LNA oligonucleotides were resuspended in nuclease-free water used to prepare 50 µM stock solutions. Huh7 cells were seeded into a 24-well plate at a density of 1×10^5^ cells per well and incubated overnight at 37°C and 5% CO_2_. The next day, the cells were transfected with LNAs at a final concentration of 400 nM LNA per well using TransIT-X2 reagent (Mirus) following the manufacturer’s protocol. For the reagent-only negative control, nuclease-free water was added instead of LNA. 4-5 hours after transfection, the cell supernatants were removed, and the cells were infected with 0.5 mL EBOV-ZsGreen inoculum at an MOI of 0.5. After incubating for 2 d at 37°C and 5% CO_2_, the cells were fixed with 10% formalin for at least 6 h and removed from the BSL-4 laboratory in accordance with approved inactivation SOPs. To stain the cell nuclei, cells were washed four times in PBS (Boston Bioproducts) and incubated with 200 mg/mL 4’,6-diamidino-2-phenylindole (DAPI, Sigma-Aldrich) for 20 min at room temperature. Images were acquired at 4x magnification on a Nikon Ti2 Eclipse microscope and Photometrics Prime BSI camera with NIS Elements AR software. Fiji Image J software (https://github.com/fiji) was used to create and merge channel images and add scale bars. Total cell quantification was performed on at least 4 images per sample using the cell detection feature in QuPath (v.0.6.0 https://qupath.github.io/) to identify DAPI-stained nuclei (73). Infection rates were quantified using the mean fluorescence signal for the green channel. Changes in infection upon LNA treatment were quantified relative to the scramble LNA treatment by ordinary one-way ANOVA for multiple comparisons.

### Synonymous mutation rate analysis

To identify evolutionary support for RNA structures by synonymous mutation rates (SMR), a multiple sequence alignment (MSA) for mammalian filoviruses was generated using the coding sequences from VP35, VP40, VP30, and VP24 of 11 mammalian filoviruses. All coding sequences were obtained from complete genomes and downloaded from NCBI using the specified accessions (**Table S4**). The codon-based alignment tool MACSE (v 2.0.7) with the default parameters was used to generate MSAs. MSAs were then visualized and edited in Jalview (v 2.10.2b1), removing stop codons and alignment columns corresponding to gaps in the reference EBOV sequence (74). SMRs at each codon position of the edited alignments were calculated in FUBAR using the default parameters (75). For each mRNA, SMR was assigned to each nucleotide in the codon triplet, and then assigned as double-stranded or single-stranded based on the EBOV mRNA structure model. SMR within individual LowSS structures were analyzed by comparing SMR between ssRNA and dsRNA by unpaired, two-tailed Student’s t-test. SMR within LowSS structures were further compared to the global CDS irrespective of base-pairing status, by ordinary one-way ANOVA for multiple comparisons.

### Covariation analysis

To identify evolutionary support for RNA structures by covariation, MSAs were generated for mammalian filoviruses using the full-length mRNA sequences for the same viruses as in SMR analysis. Full-length mRNA sequences were obtained from NCBI using the specified accessions (**Table S4**), except for the Lloviu mRNA sequences, which were determined by sequencing of the Lloviu Hungary isolate. 5’ UTR and 3’ UTR sequences were extracted from the full-length sequences and aligned on MAFFT using the L-INS-i method and default parameters (76). The 5’ UTR and 3’ UTR alignments from MAFFT and CDS alignments from MACSE were all visualized in Jalview (v 2.11.5.1) and edited to remove alignment columns corresponding to gaps in the reference EBOV sequence. The final MSAs containing the full-length mRNA sequences were generated by merging the 5’ UTR, CDS, 3’ UTR alignments in FASTA format. The full-length MSAs were then converted to Stockholm format using HMMER (v.3.4-gompi-2022b, http://hmmer.org) and the SHAPE-MaP secondary structures in dot-bracket format were manually added to the output Stockholm file using the #=GC SS_cons flag (77, 78). Identification of significantly covarying base pairs was performed using R-Scape (v 0.6.1, h470a237_4) installed from Bioconda (https://anaconda.org/bioconda/rscape/files/manage?page=2) using the –-RAFSp option (79, 80). Due to the length of the mRNAs, covariation analysis was performed in sliding windows with a 200 nt window size (--window), 100 nt sliding step (--slide), and default threshold E-value < 0.05.

## Statistical analysis

All statistical analyses and graphs were generated using GraphPad Prism 11. All statistical analyses performed were ordinary one-way ANOVA for multiple comparisons unless otherwise specified. Significance levels of comparisons are indicated in figures as ns – not significant; * – p < 0.05; ** – p < 0.01; *** – p < 0.001; **** – p < 0.0001.

## Data availability

Raw and processed chemical probing data have been deposited on the Gene Expression Omnibus repository under the accession GSE334453.

## Acknowledgements

We thank Sarah Fergione (Pyle Lab, Yale University/HHMI) for synthesizing the LNA oligos used in this study and Dr. Michael Van Zandt (New England Discovery Partners) for the kind gift of the 2A3 reagent. We also thank Mitchell R. White (Mühlberger lab, Boston University/NEIDL) for technical assistance and Dr. Robert A. Davey (Boston University/NEIDL) for providing access to imaging equipment. We are grateful to Dr. Anthony Mustoe and Lucas Kearns (Baylor College of Medicine) for guidance with fbDMS-MaP analysis. We thank Dr. Jimin Hwang, Dr. Mitchell Gulkis, Dr. Harim Jang, and Dr. Ling Xu (Pyle Lab, Yale University/HHMI) for helpful comments on the manuscript. Lastly, we thank members of the Pyle and Mühlberger labs for valuable and insightful discussions.

## Author Contributions (CRediT)

**Conceptualization**: Michelle Luo, Judith Olejnik, Kristina Meier, Tanja Hann, Elke Mühlberger, Anna Marie Pyle

**Data curation**: Michelle Luo, Judith Olejnik, Kristina Meier

**Formal analysis**: Michelle Luo, Judith Olejnik, Kristina Meier, Tanja Hann

**Funding acquisition**: Elke Mühlberger, Anna Marie Pyle **Investigation**: Michelle Luo, Judith Olejnik, Kristina Meier **Methodology**: Michelle Luo, Tanja Hann, Judith Olejnik, Kristina Meier

**Project administration**: Michelle Luo, Judith Olejnik, Kristina Meier, Tanja Hann, Elke Mühlberger, Anna Marie Pyle

**Resources**: Elke Mühlberger, Anna Marie Pyle

**Software**: Michelle Luo

**Supervision**: Michelle Luo, Judith Olejnik, Kristina Meier, Tanja Hann, Elke Mühlberger, Anna Marie Pyle

**Validation**: Michelle Luo, Judith Olejnik, Kristina Meier

**Visualization**: Michelle Luo, Judith Olejnik

**Writing – original draft**: Michelle Luo, Anna Marie Pyle

**Writing – review & editing**: Michelle Luo, Judith Olejnik, Kristina Meier, Tanja Hann, Elke Mühlberger, Anna Marie Pyle

## Supplemental Figures

**Fig S1.**
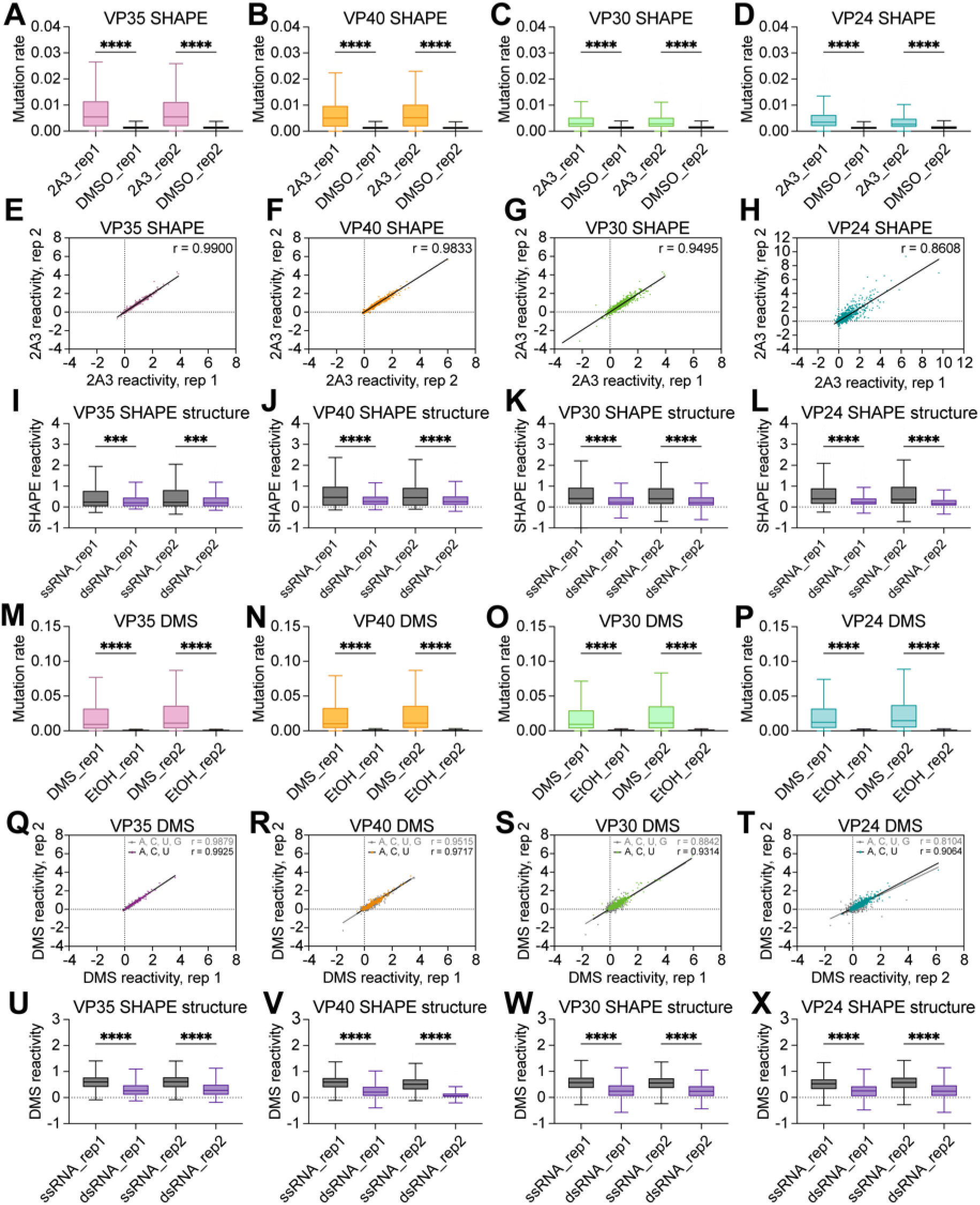
In-cell chemical probing using SHAPE-MaP and fbDMS-MaP yields high quality data for structure prediction of EBOV mRNAs. (**A-D**) Mutation rates from two independent SHAPE-MaP replicates for each mRNA. Data are shown in Tukey style with outliers omitted (boxes: interquartile range (IQR); whiskers: 1.5 x IQR; horizontal line: median). (**E-H**) Correlation of normalized SHAPE (2A3) reactivities from two independent replicates of each mRNA, shown with linear regression fit lines and Pearson correlation coefficients (r). (**I-L**) Normalized SHAPE reactivities mapped onto single– and double-stranded nucleotides of SHAPE-constrained RNA structure models, plotted as in (**A-D**). (**M-P**) Mutation rates from two independent fbDMS-MaP replicates, plotted as in (**A-D**). (**Q-T**) Correlation of normalized DMS reactivities for A/C/U nucleotides (color) or A/C/U/G nucleotides (grey) from two independent replicates for each mRNA, plotted as in (**E-H**). (**U-X**) Normalized DMS reactivities mapped onto single– and double-stranded nucleotides in the SHAPE-constrained RNA structure models, plotted as in (**I-L**). Statistical significance: *** p <0.001; **** p < 0.0001 by non-parametric Mann-Whitney U-test.

**Fig S2.**
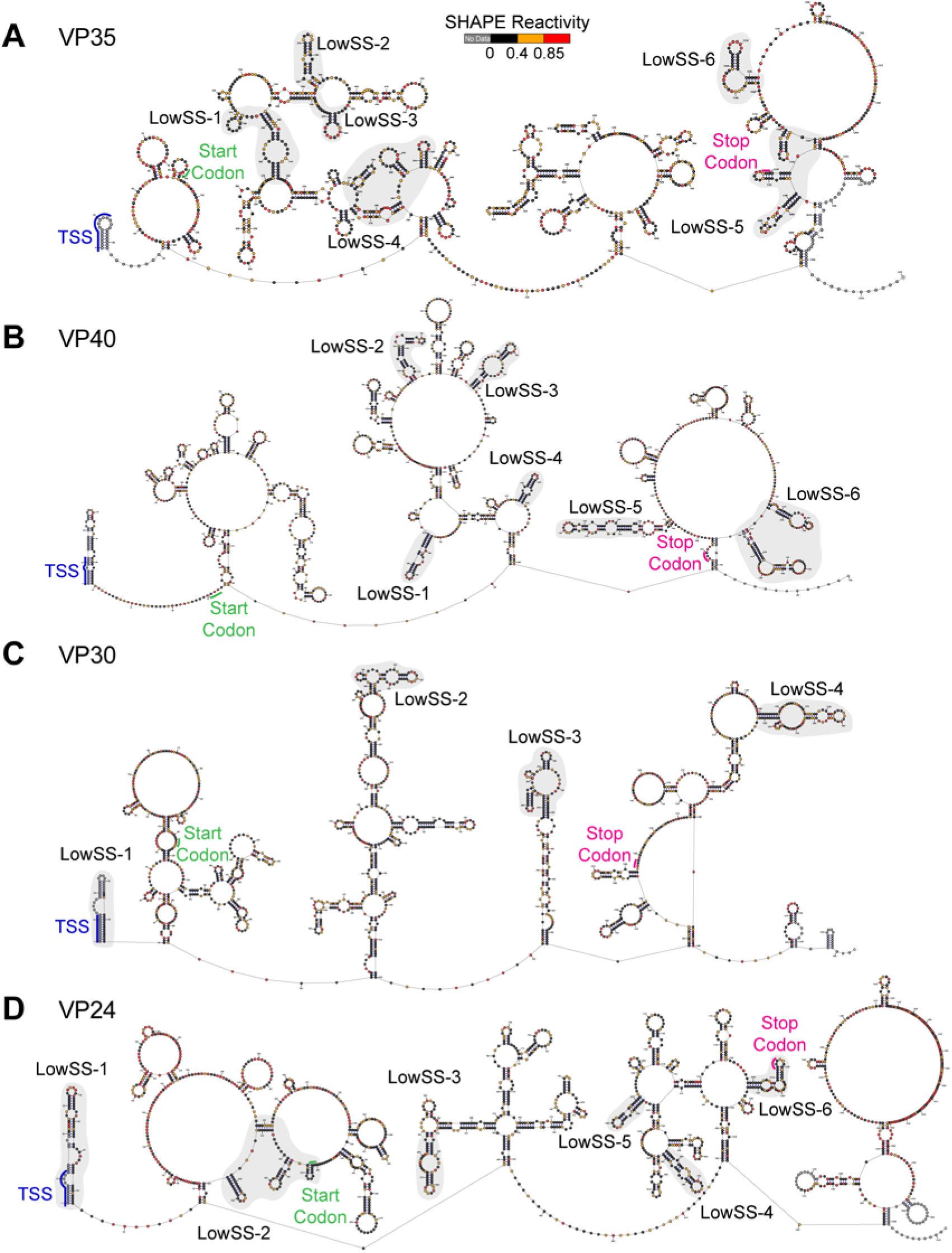
Full-length structure maps of EBOV mRNAs determined by SHAPE-MaP. (**A-D**) SHAPE-constrained structure models colored by SHAPE reactivity of (**A**) VP35 (1376 nt), (**B**) VP40 (1505 nt), (**C**) VP30 (1453 nt), (**D**) VP24 (1612 nt). LowSS regions are shaded in grey. Genetic elements: transcription start sequence (TSS, blue); start codon (green); stop codon (red).

**Fig S3.**
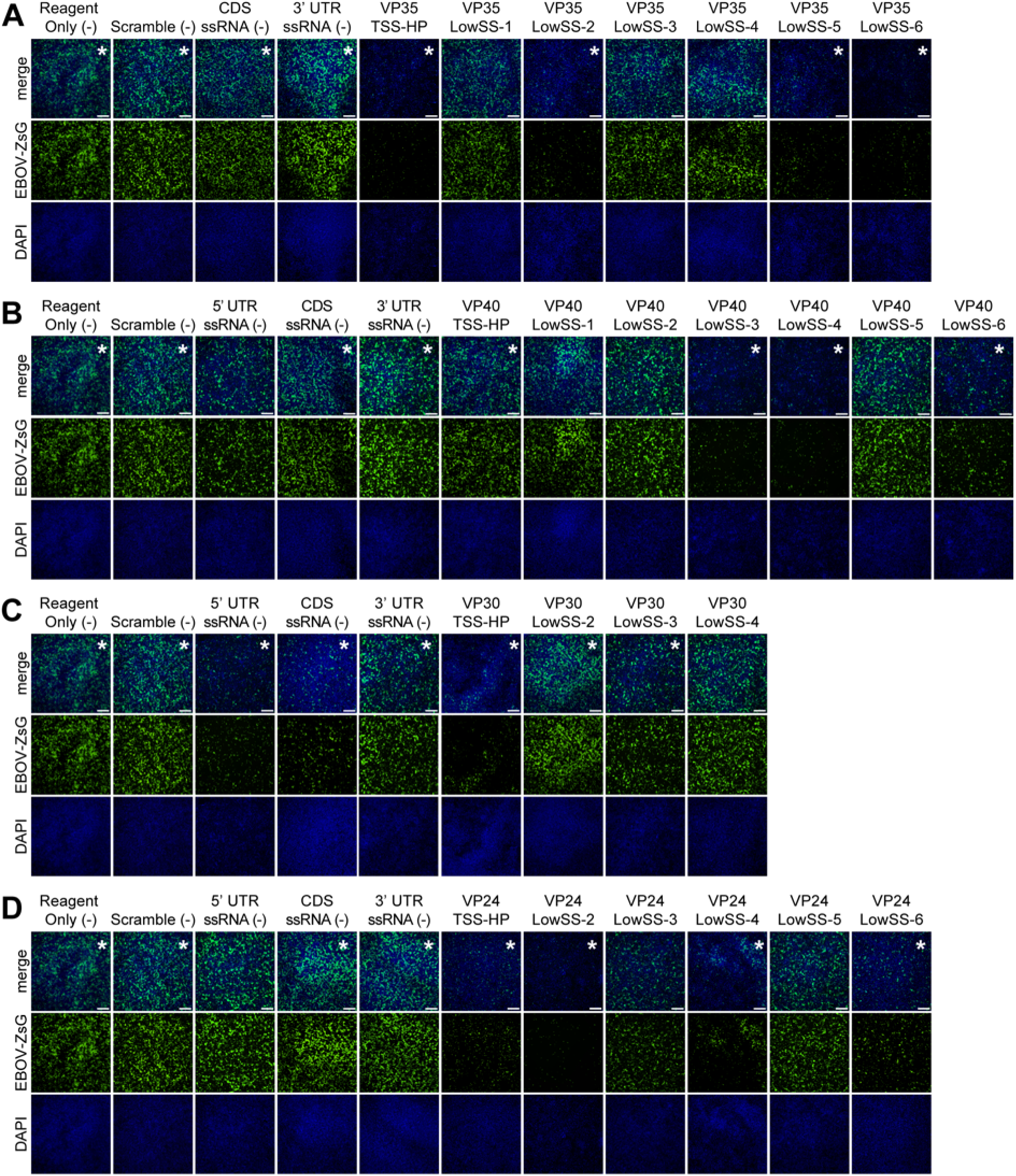
Fluorescence microscopy of EBOV-infected cells after LNA treatment. (**A-D**) Representative fluorescence microscopy of Huh7 cells after LNA targeting of RNA structures in (**A**) VP35, (**B**) VP40, (**C**) VP30, and (**D**) VP24 mRNAs. Cells were transfected with 400 nM LNA at 4-5 hours before infection (EBOV-ZsGreen-VP40; MOI = 0.5). At 2 dpi, cells were fixed with formalin and stained with DAPI. (magnification = 4x; scale bar = 500 µm). Images were selected from this set and reproduced in Fig. 3-4, denoted by an asterisk (*).

**Fig. S4.**
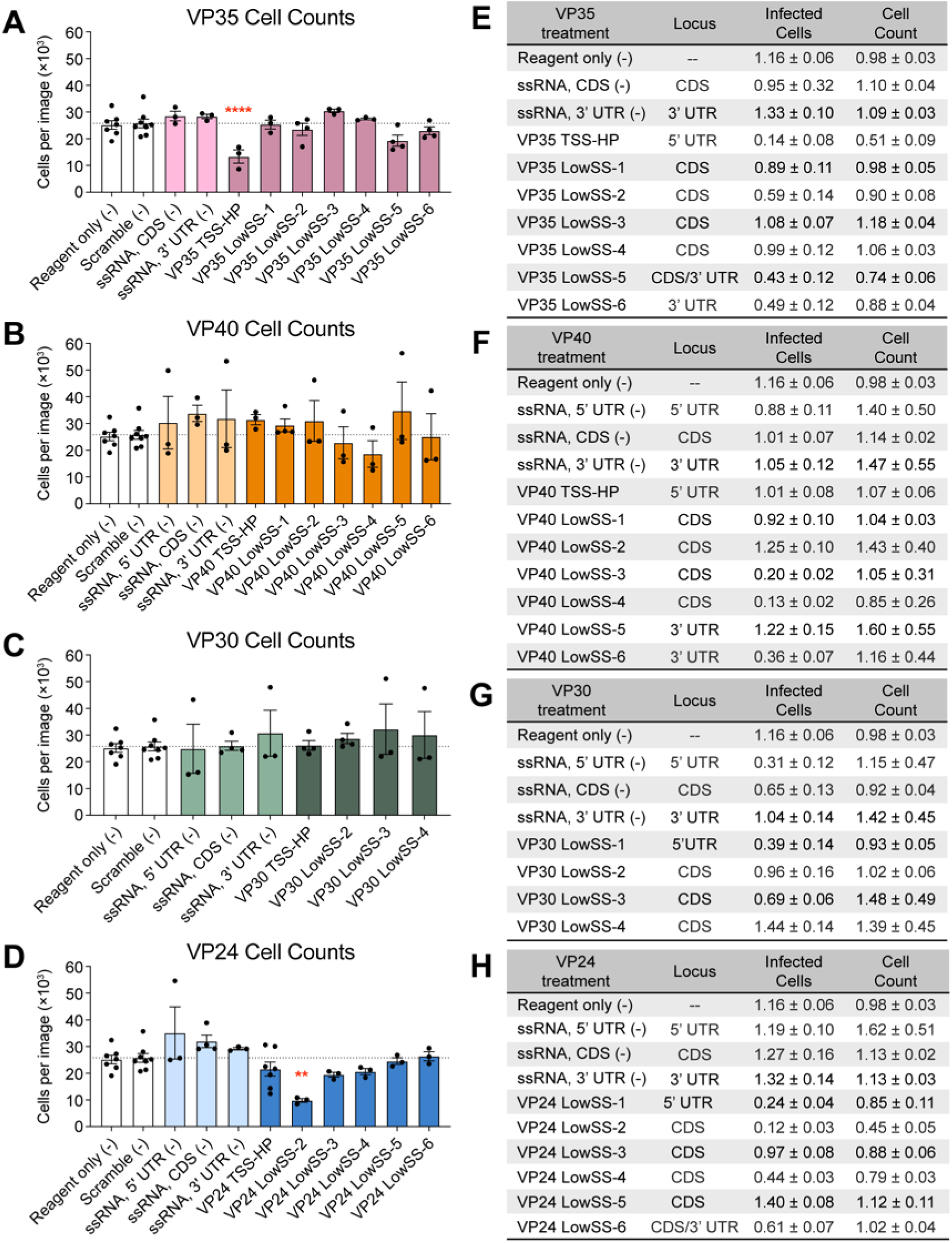
Quantification of infection and cell counts after LNA targeting. (**A-D**) Total cell counts following LNA treatment targeting LowSS structures in (**A**) VP35, (**B**) VP40, (**C**) VP30, and (**D**) VP24 mRNAs using QuPath. Data points represent the mean infection rate ± SEM from (n≥3) biological replicates (performed in technical duplicate; ≥4 images per replicate). Bars represent global negative controls (white), LNAs targeting single-stranded regions (lighter shades), and structure-disrupting LNAs (darker colors). Reagent-only and scramble negative controls are reproduced from Fig 3, as experiments were performed simultaneously with the same set of controls. TSS hairpin (TSS-HP) LNA values are reproduced from Fig 3 to compare disruption of previously predicted 5’ stem loop structures and newly identified LowSS structures. Dotted lines indicate mean cell count from scramble LNA treatment (25,214). Statistical significance relative to scramble determined by ordinary one-way ANOVA: ** p < 0.01; **** p < 0.0001. (**E-H**) Average (± SEM) infected cells or cell counts relative to scrambled LNA treatment for (**E**) VP35, (**F**) VP40, (**G**) VP30, and (**H**) VP24 mRNAs.

**Fig S5.**
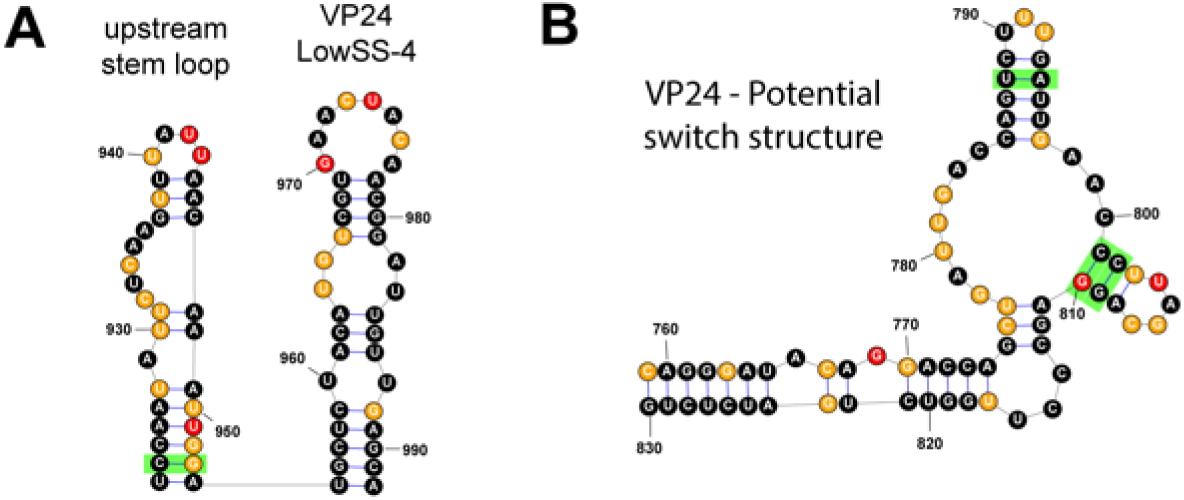
Covarying base pairs supporting structure formation in mammalian filoviruses. SHAPE-predicted structures annotated with significantly covarying base pairs (green) analyzed by R-scape (--RAFSp, Window: 200, Slide: 100, E < 0.05) for a mammalian filovirus mRNA alignment. (A) Stem loop structure upstream of VP24 LowSS-4, located in the CDS. (B) Multi-loop structure in a region with low SHAPE reactivity and high Shannon entropy in the VP24 CDS.

